# Coronavirus nucleocapsid proteins differentially antagonize RIG-I-like receptor signaling to shape pathogenicity

**DOI:** 10.64898/2026.10.09.758129

**Authors:** Yajuan Rui, Yanpu Wang, Shiqi Chen, Yan Cao, Ying Hu, Si Shen, Qiang Ding, Jiaming Su, Xiao-Fang Yu

## Abstract

Coronaviruses cause human infections with divergent clinical outcomes. Viral proteins likely play important roles in the severity of human diseases but are poorly defined. In this study, we discovered that viral nucleocapsid (N) proteins from various human coronaviruses have distinct abilities to antagonize human innate immune responses. N proteins from more pathogenic coronaviruses, such as SARS-CoV and SARS-CoV-2, have more potent activities specifically against human RIG-I-like receptor (RLR) activation than less pathogenic coronaviruses including OC43 and HKU1. Mechanistically, the SARS-CoV-2 N protein interacts with RLR pathway viral RNA sensors MDA5 and RIG-I and inhibits their interaction with MAVS. The CTD of SARS-CoV-2 N protein is an important region for RLR antagonism. Consequently, N proteins from more pathogenic coronaviruses promote viral replication more effectively than N proteins from less pathogenic coronaviruses. Blocking N protein function may be a promising antiviral strategy.

## Introduction

Coronaviruses infect humans with a striking range of clinical outcomes. Seasonal coronaviruses—HCoV-OC43, HCoV-HKU1, HCoV-229E, and HCoV-NL63—circulate endemically and typically cause mild upper respiratory illness[1, 2]. In contrast, the zoonotic coronaviruses SARS-CoV, MERS-CoV, and SARS-CoV-2 have each caused epidemic or pandemic disease with case fatality rates reaching 10%, 34%, and 2–4%, respectively[3, 4]. This divergence is not explained by receptor usage alone; all seven viruses enter human airway epithelial cells, yet only the highly pathogenic species achieve sustained replication in the lower respiratory tract and provoke dysregulated inflammatory responses[5, 6]. A central but unresolved question is which viral determinants govern this differential pathogenicity. Accumulating evidence points to the innate immune evasion capacity of individual viral proteins as a key factor: highly pathogenic coronaviruses suppress interferon (IFN) induction more effectively than seasonal counterparts, and this suppression correlates with higher viral loads and more severe disease in patients and animal models[3, 7, 8].

The first line of antiviral defense in airway epithelial cells is mounted by RIG-I-like receptors (RLRs). RIG-I and MDA5 are cytoplasmic DExD/H-box helicases that recognize distinct features of viral double-stranded RNA—RIG-I binds short 5′-triphosphorylated RNAs, while MDA5 recognizes longer dsRNA filaments formed during replication[9, 10]. Upon RNA binding, both receptors undergo conformational changes that expose N-terminal caspase activation and recruitment domains (CARDs) for K63-linked polyubiquitination by the E3 ligases TRIM25 and Riplet, enabling their interaction with the adaptor protein MAVS on mitochondrial membranes[11]. MAVS then nucleates a signaling cascade through TBK1–IRF3 and IKK–NF-κB, driving transcription of type I and type III IFNs and proinflammatory cytokines [12, 13]. MDA5 has emerged as the predominant sensor for SARS-CoV-2 in lung epithelial cells: genetic ablation of MDA5, but not RIG-I, abolishes type I and III IFN induction in response to SARS-CoV-2 infection[14, 15], and RIG-I/MDA5-mediated sensing in the airway epithelium links viral replication to macrophage-driven inflammation [16]. Recent work has further revealed that MDA5 agonists during infection are not limited to viral RNAs—MDA5 binds predominantly to aberrant cellular transcripts, including intron-containing and Alu-proximal RNAs, whose cytoplasmic accumulation reflects disruption of host RNA processing fidelity [17]. Thus, MDA5 surveys both viral replication intermediates and the integrity of cellular RNA homeostasis, positioning it as a sentinel whose activation coronavirus proteins must evade to establish infection.

RNA viruses have evolved diverse strategies to neutralize RLR signaling, and the potency of these antagonists frequently tracks with virulence. Paramyxovirus V proteins bind directly to MDA5 and RIG-I, disrupting their interaction with TRIM25 and preventing downstream MAVS activation [18, 19]. Flavivirus NS3 proteases cleave MAVS or sequester 14-3-3 proteins required for RIG-I and MDA5 signaling [20]. Picornavirus 3C and leader proteases degrade MDA5 or cleave RIG-I and MAVS to abort IFN induction [21, 22]. Coronaviruses deploy multiple proteins against the RLR pathway: the papain-like protease (PLpro) de-ISGylates MDA5 to prevent its filament assembly [23], nsp3 binds MDA5 and removes activating ubiquitin marks[24], nsp8 interferes with TRIM4-mediated K63 ubiquitination of MDA5[25], and the M protein and ORF9b disrupt RIG-I/MDA5–MAVS complexes [26, 27]. Systematic functional screens have identified the nucleocapsid (N) protein among SARS-CoV-2’s innate immune antagonists [28, 29], yet no study has compared how proteins from the same functional class—across both highly pathogenic and seasonal species—differ in their capacity to suppress RLR activation.

The nucleocapsid (N) protein is the most abundant viral protein in infected cells and performs multiple roles beyond genome packaging. N is a multidomain RNA-binding protein with an N-terminal RNA-binding domain (NTD), a C-terminal dimerization domain (CTD), and a disordered serine/arginine-rich linker (SR region)[30, 31]. Through multivalent RNA interactions, N undergoes liquid–liquid phase separation to condense the viral genome into ribonucleoprotein complexes [30, 32-34]. Several recent studies have reported that N proteins from individual coronaviruses also antagonize innate immunity. SARS-CoV-2 N blocks RIG-I ATPase activity, prevents TRIM25-mediated RIG-I ubiquitination, and suppresses STAT1/STAT2 phosphorylation to dampen both IFN induction and IFN-stimulated gene expression [35-37]. SARS-CoV-2 N also promotes MAVS SUMOylation via UBC9, attenuating MAVS ubiquitination and downstream IRF3 activation [38]. These findings establish N as a bona fide innate immune antagonist, but they have emerged piecemeal, each focused on a single virus. Whether N proteins from different human coronaviruses differ quantitatively in their anti-RLR activities—and whether such differences correlate with pathogenicity—remains unknown.

Here we show that coronavirus N proteins are differential antagonists of human RLR signaling. By comparing N proteins from human coronaviruses, we find that N proteins from highly pathogenic species (SARS-CoV, SARS-CoV-2) potently suppress RIG-I– and MDA5–MAVS signaling, whereas N proteins from seasonal coronaviruses (OC43, HKU1) exhibit markedly weaker antagonism. Mechanistically, SARS-CoV-2 N interacts directly with both MDA5 and RIG-I, blocking their engagement with MAVS. The C-terminal domain (CTD) of SARS-CoV-2 N is necessary for this RLR antagonism. Consistent with their differential immune evasion capacities, N proteins from more pathogenic coronaviruses enhance viral replication more effectively than those from seasonal strains. Our results identify N protein–RLR interaction as a determinant of coronavirus pathogenicity and suggest that disrupting this interface is a viable antiviral strategy.

## Results

### N proteins from SARS-CoV-2 and SARS-COV had more potent activities against RLR activation than N proteins from less pathogenic OC43 and HKU1

We previously reported that SARS-CoV-2 encoded N protein has the function of suppressing RLR activation [39]. It is not clear whether this is the common feature of human β coronaviruses. We therefore compared the function of various N proteins from human coronaviruses SARS-CoV, SARS-CoV-2, OC43, and HKU1 (Figure 1A). RLR pathway activation after RNA virus Sendai virus (SeV) infection results in the activation of IRF3 and NF-κB [9, 40]. The promoter region of IFNβ contains IRF3 and NF-κB binding motifs[41, 42]. Both IRF3 and NF-κB activations are required for IFNβ induction [43]. We first examined IFNβ promoter activity after SeV infection and observed a significant activation compared to uninfected controls (Figure 1B).

**Fig. 1.**
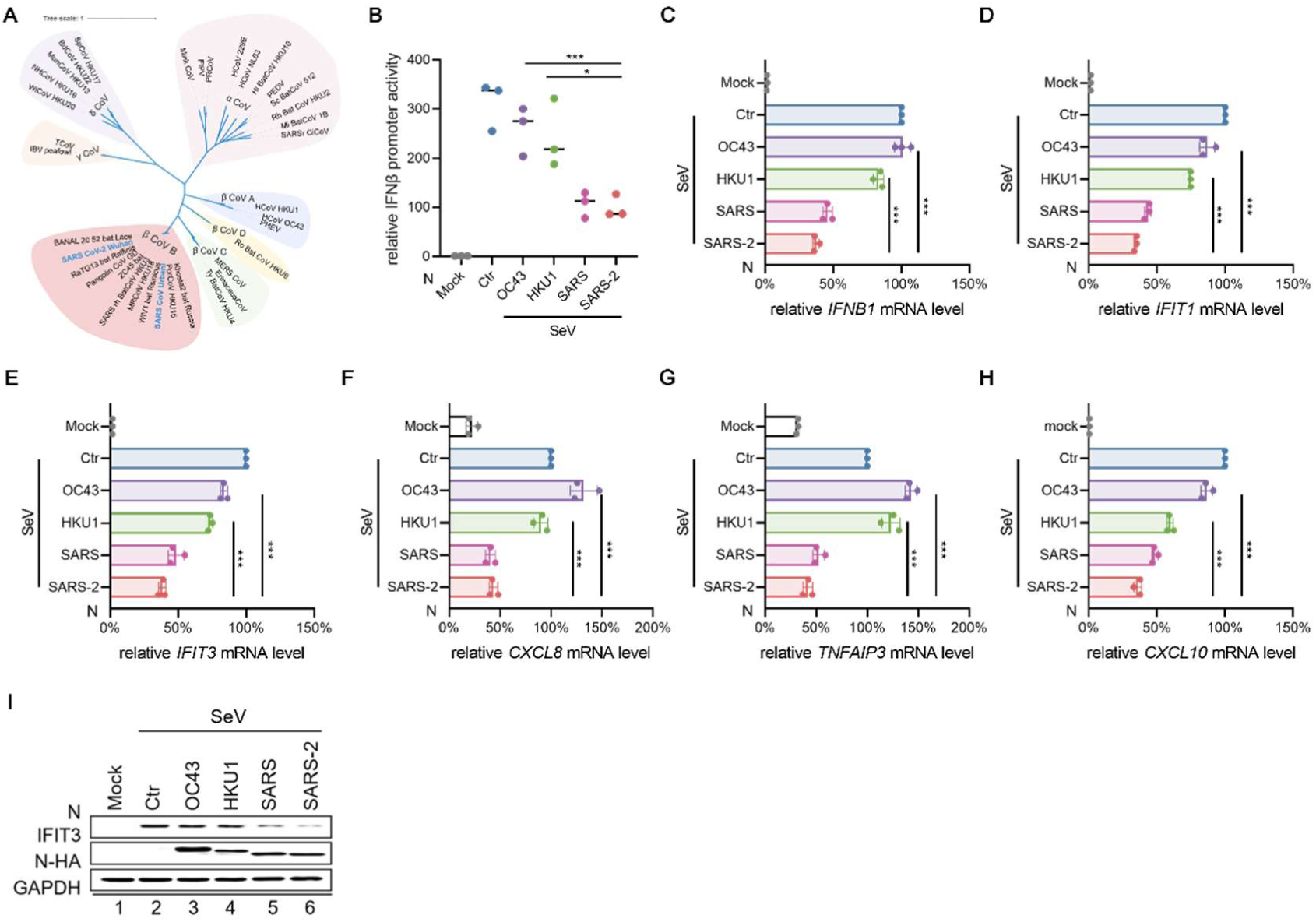
N proteins from highly pathogenic human coronaviruses potently suppress SeV-induced innate immune responses. **A,** Phylogenetic tree of N proteins from representative α-, β-, γ- and δ-coronaviruses, coloured by genus and betacoronavirus lineage: Alphacoronavirus (light pink), Betacoronavirus lineage A (light purple), lineage B (orange-red), lineage C (light green), lineage D (light yellow), Gammacoronavirus (light orange) and Deltacoronavirus (violet); SARS-CoV and SARS-CoV-2 are highlighted in bold blue. **B,** HEK293T cells were co-transfected with an IFNβ promoter–luciferase reporter, a Renilla internal control and HA-tagged N from the indicated human coronaviruses; 18 h after transfection, cells were stimulated with Sendai virus (SeV) and harvested 18 h later for dual-luciferase assay. Reporter activity in SeV-stimulated cells was set to 100%. **C–H**, Parallel cultures from b were harvested for RT–qPCR analysis of *IFNB1* (**C**), *IFIT1* (**D**), *IFIT3* (**E**), *CXCL8* (**F**), *TNFAIP3* (**G**) and *CXCL10* (**H**) mRNAs. **i**, Immunoblot analysis of HA-tagged N protein and endogenous IFIT3 in the corresponding lysates, with GAPDH as loading control. Data are mean ± s.d. of three independent biological replicates; P values were calculated by two-sided unpaired t-tests comparing the indicated N protein with SARS-CoV-2 N (***P < 0.001).

We observed that the N protein from OC43 had poor RLR suppressing activity (Figure 1B) when expressed at similar level to N protein expressed after OC43 infection (Supplementary Fig. 1A). In contrast, N proteins from SARS-CoV and SARS-CoV-2 had potent inhibitory activities toward RLR activation (Figure 1B) when expressed at comparable levels as the N protein from OC43 (Supplementary Fig. 1B). N protein from HKU1 also had impaired abilities to suppress SeV-induced IFNβ promoter activation (Figure 1B). Similar results were obtained in SeV-induced NF-κB, IRF3 and ISRE reporter assays (Supplementary Fig. 1C-E). We next examined IFNβ mRNA expression and N proteins from SARS-CoV and SARS-CoV-2 inhibited SeV infection induced IFNβ mRNA upregulation more effectively than N proteins from OC43 and HKU1 (Figure 1C).

RLR activation induced IRF3 and NF-κB signalling to generate type-I interferon and downstream interferon-stimulated genes (ISGs) [44, 45]. NF-κB signalling could also produce NF-κB stimulated genes (NSGs) [46, 47]. RLR triggered ISGs (*IFIT1* and *IFIT3*) mRNA upregulation (Figure 1D-E) and NF-κB triggered *CXCL8*, *TNFAIP3*, *CXCL10* activation (Figure 1F-H) were also effectively suppressed by N proteins from SARS-CoV and SARS-CoV-2. N proteins from OC43 and HKU1 had reduced abilities to suppress RLR triggered ISGs (*IFIT1* and *IFIT3*) mRNA upregulation (Figure 1D-E) and NF-κB triggered *CXCL8*, *TNFAIP3*, *CXCL10* activation (Figure 1F-H). IFIT3 protein levels were also examined and N proteins from SARS-CoV and SARS-CoV-2 reduced IFIT3 expression more effectively than N proteins from OC43 and HKU1 after RLR activation (Figure 1I), consistent with the mRNA results (Figure 1E). Similar to SeV infection triggered RLR activation, poly(I:C) triggered RLR activation was also suppressed by N proteins from SARS-CoV and SARS-CoV-2 (Supplementary Fig. 1F-G). N proteins from OC43 and HKU1 had impaired function suppressing poly(I:C) triggered RLR activation (Supplementary Fig. 1F-G).

### Selectively targeting RLR pathways factors by SARS-CoV-2 N protein

To delineate the molecular basis and examine the mechanism of SARS-CoV-2 N protein mediated suppression of RLR pathway, we systematically evaluated N protein interference at successive tiers of the RLR signaling pathway (Figure 2A). Individual components of the pathway including the cytosolic RNA sensors RIG-I and MDA5, the adaptor MAVS, the downstream kinases TBK1, IKKα, IKKβ, and IKKε, and the transcription factor NF-κB p65 were ectopically expressed in the presence or absence of N protein, and downstream signaling output was quantified. All these factors have the ability to activate NF-κB signalling [48-52]. We observed potent inhibition of MDA5 (Figure 2B) and RIG-I (Figure 2C) triggered NF-κB activation by SARS-CoV2 N protein. In contrast, SARS-CoV-2 N protein had little effect on IKKβ (Figure 2D), TBK1 (Figure 2E) or p65 (Figure 2F) triggered NF-κB activation. A moderate inhibition of MAVS triggered NF-κB activation by SARS-CoV-2 N protein was observed (Figure 2G). Consistent with our previous report [39], SARS-CoV-2 N protein had no effect on cGAS-STING triggered NF-κB activation (Figure 2H). SARS-CoV-2 N protein had only minor effect on MYD88 triggered NF-κB activation compared to its effect on SeV infection triggered NF-κB activation (Figure 2H). RLR, cGAS-STING, and TRIF activation could also induce IFNβ activation [53]. We detected efficient inhibition of SeV infection triggered IFNβ activation but not cGAS-STING or TRIF triggered IFNβ activation by SARS-CoV-2 N protein (Figure 2I). These data revealed a selective inhibition of RLR but not cGAS-STING or TLR hub protein function by SARS-CoV-2 N protein.

**Fig. 2.**
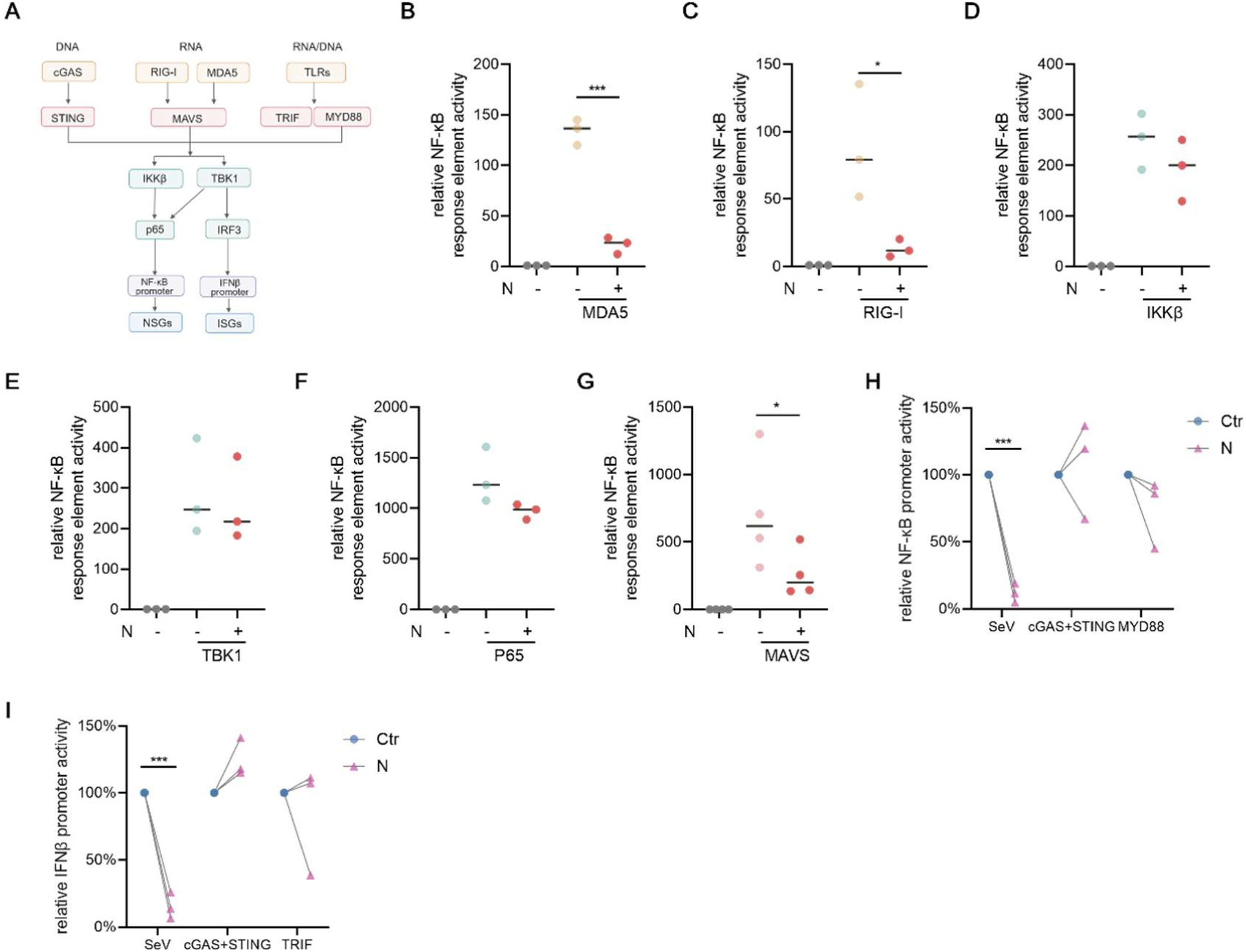
The SARS-CoV-2 N protein selectivly inhibits RLR signalling. **A,** Schematic of the RLR, TLR and cGAS–STING innate immune signalling pathways. **B–G,** HEK293T cells were co-transfected with an NF-κB response element–luciferase reporter, a Renilla internal control, and HA-tagged SARS-CoV-2 N (S2-N-HA) together with MDA5 (**B**), RIG-I (**C**), IKKβ (**D**), TBK1 (**E**), p65 (**F**) or MAVS (**G**); cells were harvested 36 h after transfection for dual-luciferase assay. **H,I,** HEK293T cells were co-transfected with NF-κB response element (**H**) or an IFNβ promoter (**I**) reporter, a Renilla internal control, and cGAS plus STING, or TRIF, or MYD88, with or without S2-N-HA, and harvested 36 h after transfection; where indicated, SeV was added 18 h after transfection and cells were harvested 18 h later. Reporter activity in cells expressing the indicated activator alone was set to 100%. Data are mean ± s.d. of three independent biological replicates; P values were calculated by two-sided unpaired t-tests comparing each condition with the corresponding condition lacking S2-N-HA (***P < 0.001, *P < 0.05).

### SARS-CoV-2 N protein targets RNA sensor proteins in the RLR pathway

To examine the mechanism of SARS-CoV-2 N protein mediated suppression of RLR pathway, we evaluated its interaction with key pathway factors. We discovered that SARS-CoV-2 N protein interacted with MDA5, MAVS, and RIG-I but had no detectable interaction with IKKα, IKKβ, IKKε, or TBK1 (Figure 3A), consistent with the results that SARS-CoV-2 N protein did not interfere with IKKβ, or TBK1 function (Figure 2D-E). N protein of SARS-CoV-2 also did not interact with STING, TRIF, or MYD88 molecules (Figure 3A). MDA5 and RIG-I recognize target RNA and subsequently use their exposed CARD domains to associate with MAVS and trigger innate immune activation [9, 11, 48]. The CARD domains of MDA5 and RIG-I are important for MAVS binding and activation [54-56]. We observed that SARS-CoV-2 N protein could also interact with MDA5 CARD domain (Figure 3A). MDA5 CARD domain triggered NF-κB activation was blocked by SARS-CoV-2 N protein (Figure 3B). We also observed that SARS-CoV-2 N protein blocked the interaction between MDA5 CARD domain with MAVS (Figure 3C). Similarly, RIG-I CARD domain triggered NF-κB activation was blocked by SARS-CoV-2 N protein (Supplementary Fig. 3A). The interaction between RIG-I CARD domain with MAVS was also blocked by SARS-CoV-2 N protein (Supplementary Fig. 3B).

**Fig. 3.**
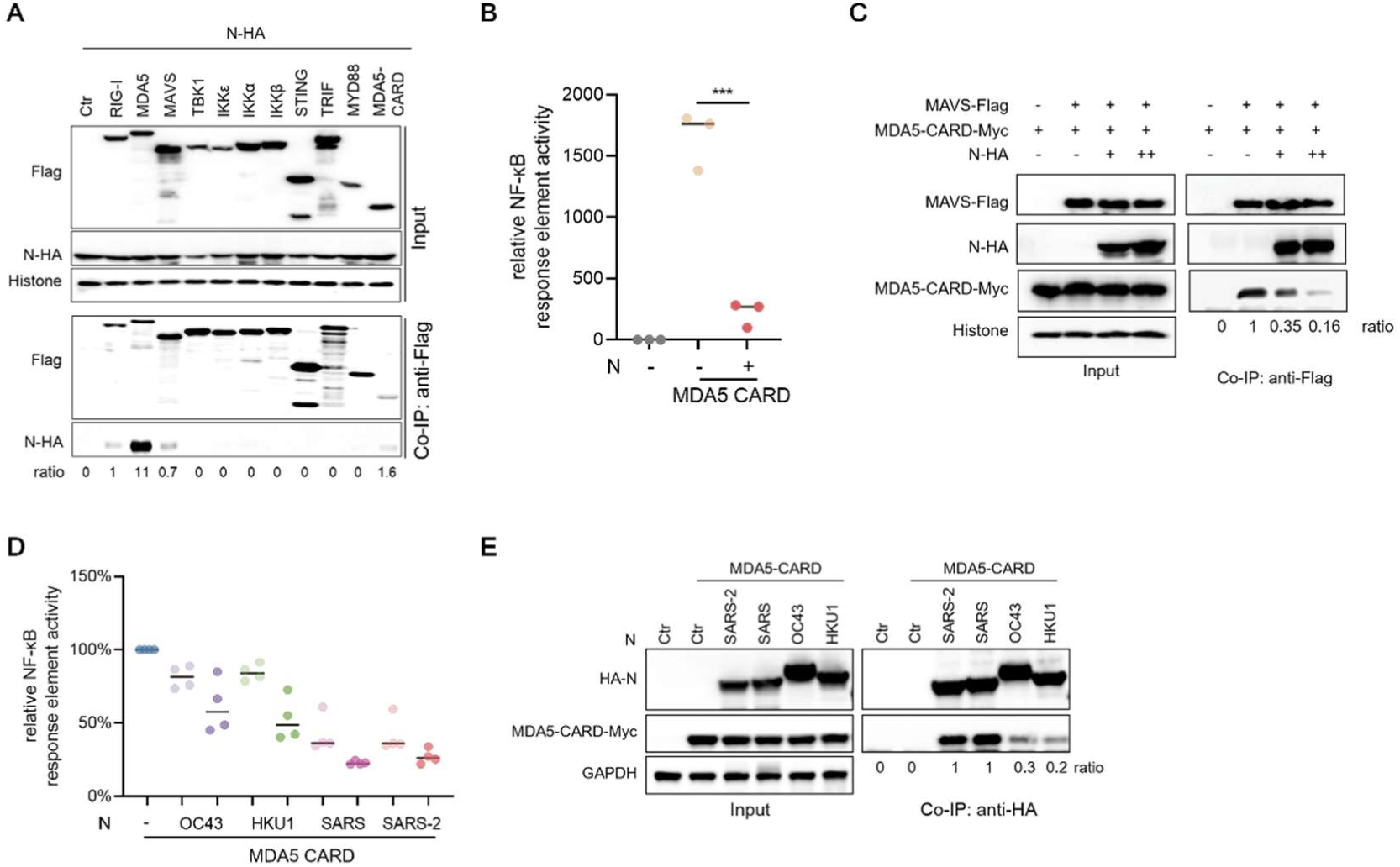
The SARS-CoV-2 N protein interacts with the CARD domains of MDA5 and disrupts the MDA5–MAVS interaction. **A,** HEK293T cells were co-transfected with S2-N-HA and Flag-tagged innate immune signalling molecules as indicated; 24 h later, cell lysates were subjected to anti-Flag immunoprecipitation (IP) followed by immunoblot analysis of co-precipitated N-HA, with input lysates shown. **B,** HEK293T cells were co-transfected with an NF-κB response element reporter, a Renilla internal control, MDA5 CARD and increasing doses of S2-N-HA as indicated, and harvested 36 h after transfection. **C,** HEK293T cells were co-transfected with MAVS-Flag, MDA5 CARD-Myc and increasing doses of S2-N-HA as indicated; 36 h later, cell lysates were subjected to anti-Flag IP followed by immunoblot analysis of co-precipitated MDA5 CARD-Myc. **D,** As in **B**, but with increasing doses of HA-tagged N from the indicated human coronaviruses of high and low pathogenicity. **E,** HEK293T cells were co-transfected with MDA5 CARD-Myc and HA-tagged N from the indicated viruses; 36 h later, cell lysates were subjected to anti-HA IP followed by immunoblot analysis of co-precipitated MDA5 CARD-Myc. Reporter data are mean ± s.d. of three independent biological replicates; P values were calculated by two-sided unpaired t-tests comparing each condition with MDA5 CARD alone (***P < 0.001).

We have detected N proteins from pathogenic coronaviruses blocked SeV infection triggered innate immune activation more effectively than N proteins from less pathogenic coronaviruses (Figure 1). We therefore compared their abilities to suppress MDA5 CARD domain triggered NF-κB activation. Again, N proteins from pathogenic coronaviruses (SARS-CoV and SARS-CoV-2) blocked MDA5 CARD domain triggered innate immune activation (Figure 3D, Supplementary Fig. 3C) more effectively than N proteins from less pathogenic coronaviruses (OC43 and HKU1). N proteins from SARS-CoV and SARS-CoV-2 interacted with MDA5 CARD domain more efficiently than N proteins from OC43 and HKU1 (Figure 3E).

### The CTD domain of SARS-CoV-2 N protein is critical for MDA5 interaction and RLR pathway inhibition

The N proteins from pathogenic human coronaviruses (SARS-CoV and SARS-CoV-2) were more effective in RLR pathway inhibition than those from less pathogenic human coronaviruses (OC43 and HKU1). To map the functional domain of N protein involved in RLR pathway inhibition, we generated a panel of truncation mutants of SARS-CoV-2 N proteins (Figure 4A). The C-terminal intrinsic disorder domain which contains nuclear targeting signal [57, 58] was found to be dispensable for RLR inhibition (Figure 4B, Supplementary Fig. 4A). In contrast, the CTD or the protein-protein interaction domain was essential for N protein mediated RLR inhibition (Figure 4B, Supplementary Fig. 4A). The CTD domain was also important for N protein mediated IRF3 phosphorylation inhibition after SeV infection (Figure 4C). The abilities of full-length and various truncation mutants of SARS-CoV-2 N proteins to interact with MDA5 correlated well with their abilities to inhibit RLR activation (Figure 4D, Supplementary Fig. 4B-C). The CTD domain appeared to be critical for SARS-CoV-2 N protein and MDA5 interaction (Figure 4D).

**Fig. 4.**
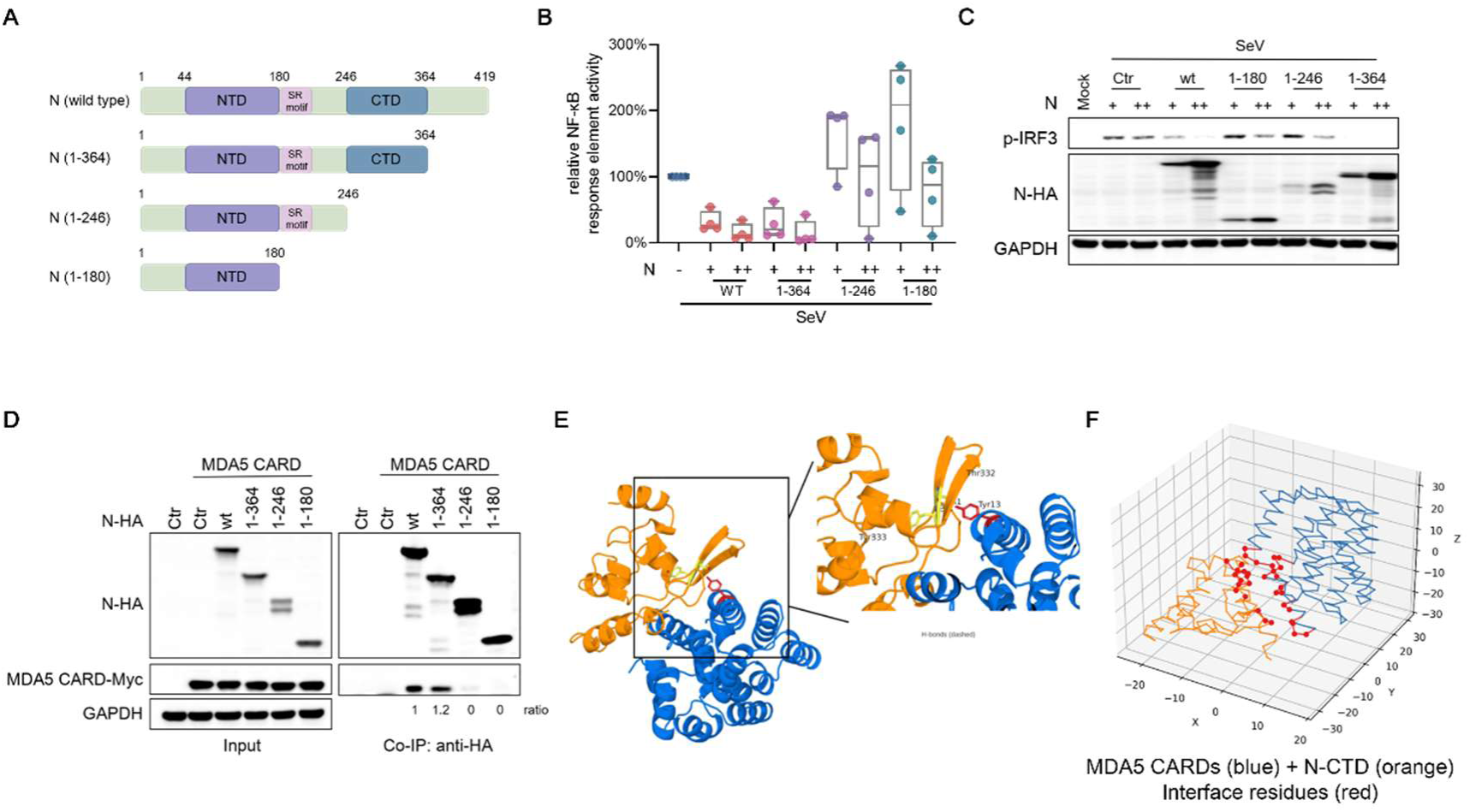
The CTD of the SARS-CoV-2 N protein is critical for MDA5 interaction and RLR pathway inhibition. **A,** Schematic of SARS-CoV-2 N protein domain organization (N-arm, NTD, SR-rich linker region, CTD, C-tail) and the truncation mutants used in this study. **B,** HEK293T cells were co-transfected with an NF-κB response element reporter, a Renilla internal control and the indicated HA-tagged N truncation mutants; 18 h after transfection, cells were stimulated with SeV and harvested 18 h later. Reporter activity in SeV-stimulated cells was set to 100%. **C,** Immunoblot analysis of IRF3 phosphorylation and N-HA expression in lysates from the corresponding samples, with GAPDH as loading control. **D,** HEK293T cells were co-transfected with MDA5 CARD-Myc and the indicated HA-tagged N truncation mutants; 24 h later, cell lysates were subjected to anti-HA IP followed by immunoblot analysis of co-precipitated MDA5 CARD-Myc. **E,** Boltz-2-predicted complex of the SARS-CoV-2 N CTD (residues 247–365; orange) and the tandem CARDs of human MDA5 (residues 1–200; blue). Left, overall docking configuration; right, close-up of the predicted interface, in which MDA5 Tyr13 forms hydrogen bonds (dashed lines) with N-CTD Thr332 and Tyr333. **F,** Interface contact map of the Boltz-2-predicted complex, showing hydrogen bonds (<3.5 Å) and hydrophobic contacts (<5.0 Å) between the 19 MDA5 and 15 N-CTD interface residues; no salt bridges were detected. Reporter data are mean ± s.d. of three independent biological replicates;

To gain structural insight into how the CTD engages MDA5 CARD domain, we used the Boltz-2 co-folding model [59, 60] to predict the complex between the SARS-CoV-2 N CTD (residues 247–365) and the tandem CARDs of human MDA5 (residues 1–200) (Figure 4E). Both chains were folded with high confidence (pLDDT 0.90), whereas the predicted interface showed a modest ipTM (0.15; Methods), a signature of the transient, low-affinity interactions typical of viral immune antagonists [18, 61]. The N-CTD engaged a bipartite surface on MDA5 comprising the CARD1 amino-terminal helix (residues 1–21) and the CARD1–CARD2 linker (residues 99–106).

This surface placement was independently reproduced by the Chai-1 co-folding model: in all five Chai-1 predictions, the N-CTD contacted the same bipartite MDA5 surface, with no contacts elsewhere on MDA5, and MDA5 Tyr13 lay at the contact hub in every model. Notably, this surface overlaps extensively with the MDA5 surface that recruits the MAVS CARD in the cryo-EM structure of the active signalling complex (PDB 7DNI), and MDA5 Tyr13 — which forms two hydrogen bonds with N-CTD Thr332 and Tyr333 in the Boltz-2 model (2.07 and 3.40 Å) — also anchors MAVS binding. Interface analysis of the Boltz-2 model identified 19 MDA5 and 15 N-CTD contact residues, connected by seven hydrogen bonds and six hydrophobic contacts but no salt bridges (Figure 4F) — a profile dominated by polar and hydrophobic contacts rather than strong electrostatics, again consistent with a transient, reversible interaction. Whereas the MDA5-facing surface was robust across methods, the precise N-CTD contact patch differed between Boltz-2 (centred on residues 332–338) and Chai-1 (shifted to residues 338–364), both within the CTD dimerization-region lobe, as expected for the binding orientation of a transient interaction (Methods). Together with the truncation analysis, these results support a model in which the N-CTD directly occupies the MAVS-recruitment surface of MDA5, competitively blocking MDA5–MAVS signalling.

### Divergent abilities of coronavirus N proteins for the promotion of viral replication in target cells

RLR activation triggers antiviral function[40]. N proteins of coronaviruses play important roles during viral assembly in virus producing cells [62, 63]. We wondered whether N proteins in the target cells could promote viral infection (Figure 5A). Indeed, we observed that N proteins of SARS-CoV-2 in virus infection target cells significantly promoted viral replication as monitored by the production of viral spike proteins or the GFP protein (Figure 5B-C, Supplementary Fig. 5) in SARS-CoV-2 GFP/ΔN replicon (trVLP) system[64].

**Fig. 5.**
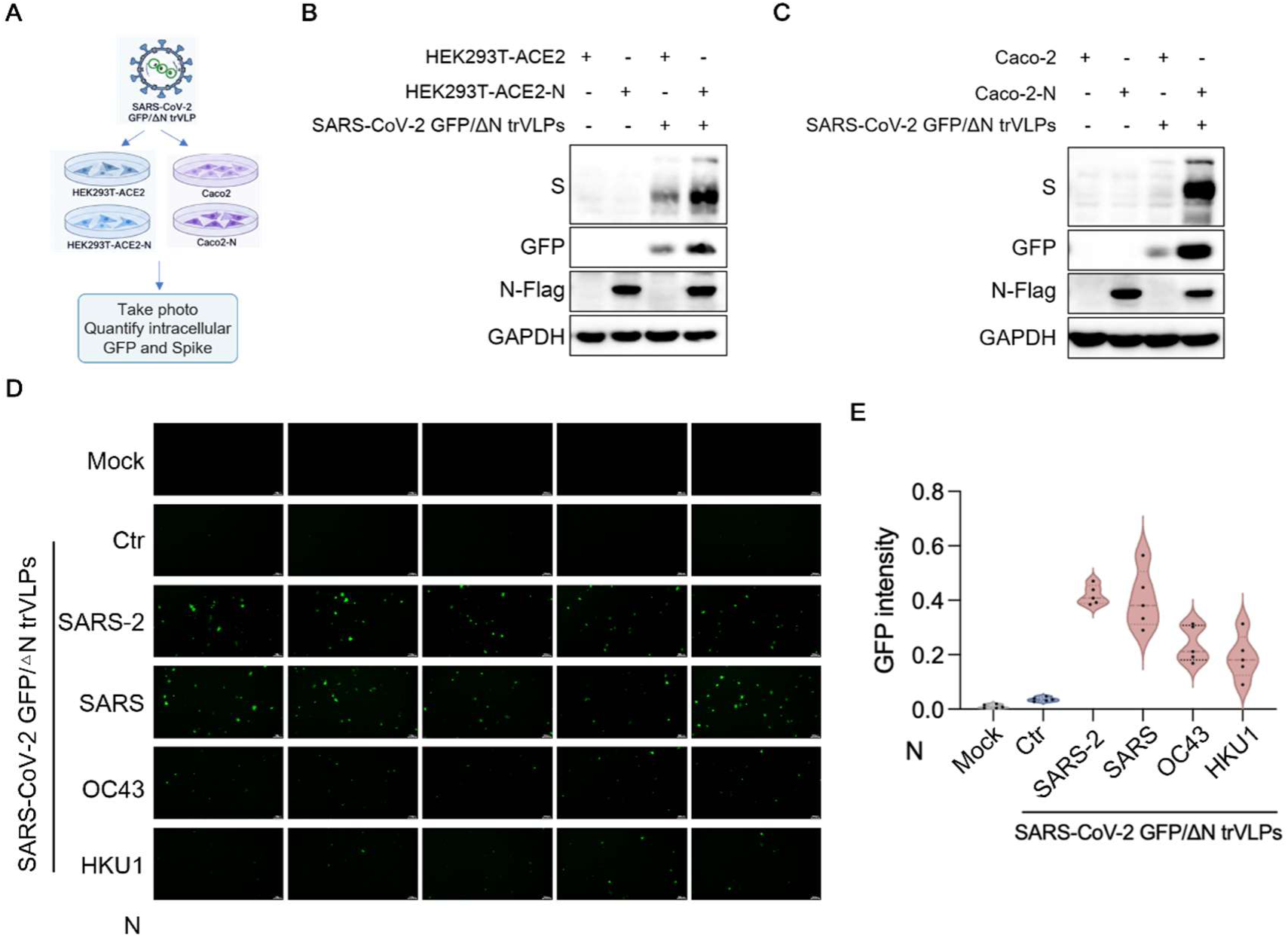
Coronavirus N protein expression enhances SARS-CoV-2 replicon replication. **A,** Schematic of the experimental workflow. **B,** HEK293T-ACE2-Flag and HEK293T-ACE2-N stable cell lines were infected with a SARS-CoV-2 replicon; 24 h later, cell lysates were immunoblotted for viral Spike protein and replicon-encoded GFP, with GAPDH as loading control. **C,** As in **B**, using Caco-2 and Caco-2-N stable cell lines. **D,** HEK293T-ACE2-Flag cells were transfected with HA-tagged N from the indicated viruses; 18 h later, cells were infected with SARS-CoV-2 replicon and imaged for GFP fluorescence 24 h after infection. **E,** Quantification of GFP fluorescence intensity from **d** using ImageJ. Data are mean ± s.d. of three independent biological replicates;

Coronaviruses N proteins displayed variable activities against RLR activation. We thus evaluated viral replication promotions of different N proteins. We observed that N proteins of SARS-CoV and SARS-CoV-2 significantly promoted SARS-CoV-2 replicon replication (Figure 5D-E). On the other hand, N proteins from OC43, HKU1 had impaired abilities to promote viral replication (Figure 5D-E). The abilities to enhance viral replication by various N proteins correlated well with their abilities to suppress RLR pathway activation (Figure 1B)

We have identified the CTD domain of SARS-CoV-2 N protein as an important region for RLR antagonism (Figure 4). It is not clear which region of SARS-CoV-2 N protein contributes to viral promotion function. Similar to RLR antagonism, the CTD region was found to be important for the promotion of SARS-CoV-2 replication (Figure 6A-B, Supplementary Fig. 6A). The C-terminal IDR domain appeared to be dispensable for viral promotion (Figure 6A). Again, viral promotion function correlated well with N protein’s RLR antagonist activities for SARS-CoV-2 N protein truncation mutants (Figure 4). Viral replication of unrelated RNA viruses (VSV) was also enhanced by N proteins of SARS-CoV-2 in target cells (Figure 6C-E).

**Fig. 6.**
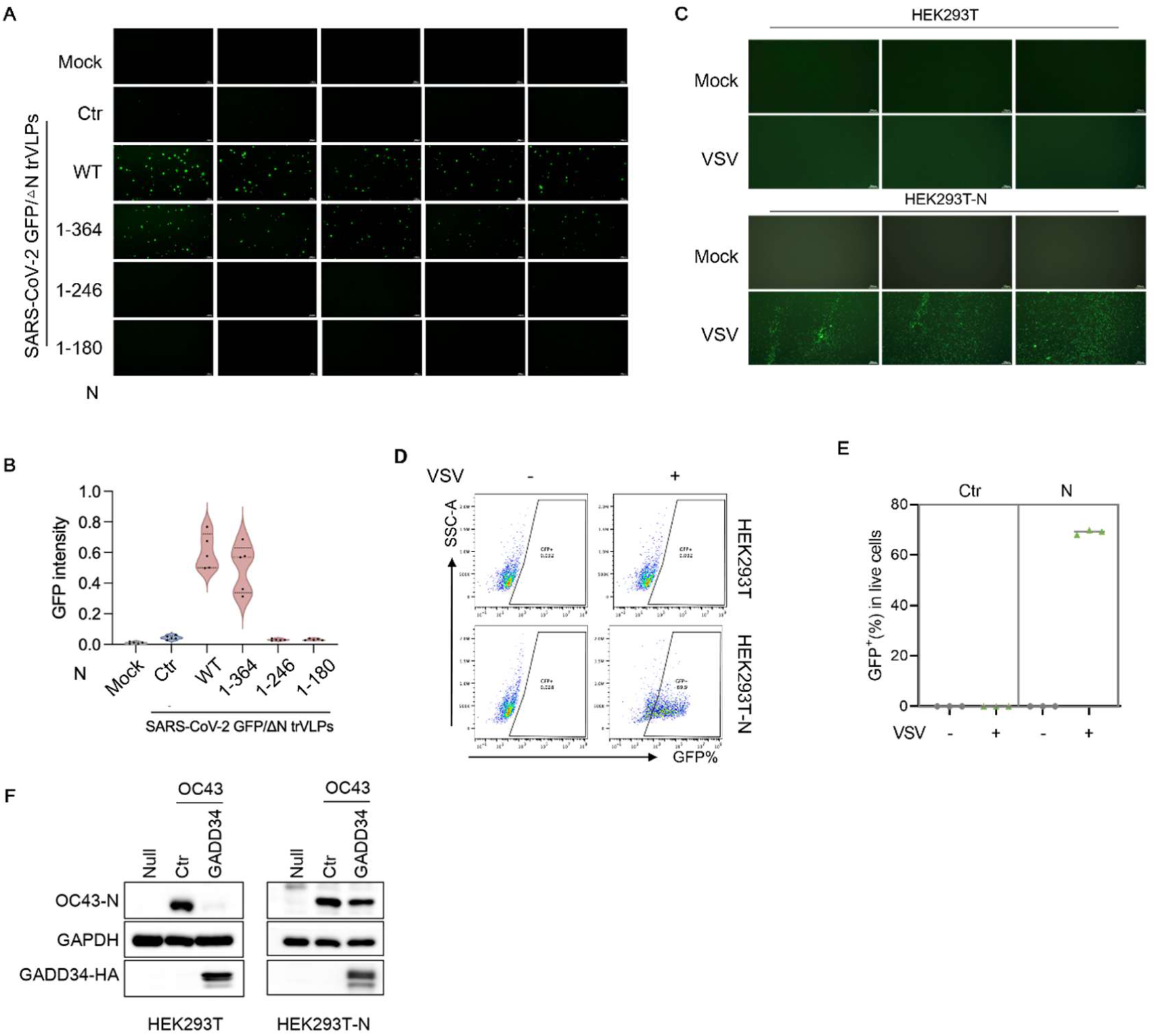
The CTD of N protein promotes VSV replication. **A,** HEK293T-ACE2 cells were transfected with the indicated HA-tagged N truncation mutants; 18 h later, cells were infected with SARS-CoV-2 replicon and imaged for GFP fluorescence 24 h after infection. **B,** Quantification of GFP fluorescence intensity from **A** using ImageJ. **c,** HEK293T and HEK293T-N-HA stable cell lines were infected with VSV-GFP; 15 h later, cells were imaged for GFP fluorescence. **D,** Flow-cytometric analysis of GFP intensity in cells from **C**. **E,** Quantification of **D**. **F,** HEK293T and HEK293T-N-HA stable cell lines were transfected with HA-GADD34; 18 h later, cells were infected with HCoV-OC43 and harvested 18 h after infection for immunoblot analysis of OC43 N protein and HA-GADD34 expression, with GAPDH as loading control. Data are mean ± s.d. of three independent biological replicates.

We have previously shown that GADD34 is a restriction factor against SARS-CoV-2 (47). Since SARS-CoV-2 N protein can enhance SARS-CoV-2 infection, we examined the ability of N protein for GADD34 antagonism. Indeed, N protein of SARS-CoV-2 suppressed the antiviral function of GADD34 (Figure 6F).

## Discussion

The structural protein N plays important roles during human coronavirus replication including viral genome packaging, virus assembly, and early steps of viral infection[30, 62, 63]. N proteins of SARS-CoV-2 and SARS-CoV have been reported to inhibit human innate immune responses[30, 35, 36]. Whether N proteins from diverse human coronaviruses have similar abilities to antagonize host innate immune systems are not well studied. We discovered that N proteins from SARS-CoV-2 and SARS-CoV exhibited more potent functions against human RLR activation than OC43 and HKU1 (Figure 1). Our results are consistent with the observation that pathogenic human coronaviruses suppressed innate immune activation more effectively than less pathogenic human coronaviruses [65]. These results suggest that antagonism of RLR-mediated antiviral signaling is a functionally divergent property of coronavirus N proteins. The potency of this antagonism may be linked to viral replication efficiency and, possibly, pathogenic potential in humans. The observation that N proteins from SARS-CoV and SARS-CoV-2, two highly pathogenic zoonotic coronaviruses, exert stronger suppression of RLR activation than those from the endemic, generally milder OC43 and HKU1 beta coronaviruses raises the possibility that enhanced innate immune evasion by N contributed to the severity of SARS and COVID-19. Conceivably, robust N-mediated antagonism of RLR signaling during early infection could delay or dampen type I interferon responses, thereby allowing high-level viral replication before adaptive immunity is engaged, a scenario consistent with the exaggerated viral loads and dysregulated interferon responses observed in severe COVID-19 patients [66-68].

Systematic evaluation of key factors in the RLR pathway identified RNA sensor MDA5 and RIG-I as the more important targets of SARS-CoV-2 N protein (Figure 2). N protein of SARS-CoV-2 preferentially binds MDA5 and RIG-I (Figure 3A). Signalling mediated by MDA5 and RIG-I but not downstream factors including TBK1, IKKβ, or p65 was selectively depressed by SARS-CoV-2 N proteins (Figure 2B-G). This selective inhibition pattern is biologically meaningful since by acting on the most upstream components of the pathway, N protein can block RLR signaling at its inception, before signal amplification through MAVS and the kinase cascade occurs, achieving efficient immunosuppression with a single molecular interaction. The apparent evolutionary choice of targeting the most upstream key factors in the RLR pathway as antagonism targets may therefore reflect a strategy of potent but pathway-specific immune evasion that minimizes global changes to the host cells which may be detrimental to viral replication.

Targeting both MDA5 and RIG-I allows SARS-CoV-2 N proteins to intercept the two principal modes of coronavirus RNA detection. RIG-I recognizes short 5′-triphosphate/diphosphate-bearing RNAs, whereas MDA5 senses long double-stranded RNA replication intermediates [69, 70]. Targeting MDA5 is particularly relevant given that coronavirus replication generates abundant dsRNA [54, 71, 72]. Dual targeting thus ensures that neither route of viral RNA sensing can effectively trigger the innate immune responses during viral infection. It is also notable that N protein binds viral genomic RNA with high avidity; association of N with RIG-I and MDA5 may simultaneously shield viral RNA from sensor access [71].

After binding to target RNA, MDA5 undergoes conformational changes and exposes the CARD domain [8, 10]. CARD domain of MDA5 binds MAVS and triggers MAVS LLPS formation to induce downstream signalling [55, 56]. Our study showed that SARS-CoV-2 N protein could associate with CARD domain of MDA5 and inhibit its interaction with MAVS and signaling function (Figure 3C). In contrast, N proteins of OC43 and HKU1 have impaired ability to bind MDA5 CARD domain. These results indicate that recognition of MDA5 CARD domain is important for N protein mediated RLR pathway inhibition. Furthermore, we discovered that the CTD domain of SARS-CoV-2 N protein was important for MDA5 CARD domain interaction (Figure 4D). Collectively, our data revealed an evolutionarily acquired function by more pathogenic SARS-COV2 N protein than N proteins from less pathogenic coronaviruses OC43 and HKU1.

Consistent with the observation that RLR downstream factors such as TBK1, IKKα, IKKβ, IKKε, or p65 were not targeted by N protein of SARS-CoV-2, we did not detect suppression of cGAS-STING activation by SARS-COV2 N proteins (Figure 2H). Signalling mediated by TRIF and MYD88 which are key hub proteins of the TLR pathway were also not significantly inhibited by the N proteins of SARS-CoV-2 (Figure 2H-I). Future experiments will be required to determine whether TLR molecules could be affected by N proteins of coronaviruses.

It is well established that N proteins of coronaviruses play essential roles in virus-producing cells [62, 63]. N proteins are one of the most abundant viral proteins in the released virions[73]. After viral entry, N proteins are released into target cells. We observed that N proteins in target cells can promote early viral infection. N proteins from SARS-CoV and SARS-CoV-2 could promote viral infection in target cells. In contrast, N proteins from OC43 and HKU1 were defective for such function. RLR activation could suppress coronavirus infection (Supplementary Fig. 6B). One possibility is that SARS-CoV-2 N protein acts directly on virus-targeted cells to suppress the induction of antiviral effectors downstream of RLR activation, thereby creating a window of opportunity for incoming viral genomes to replicate. Upon entry into a permissive cell, the viral RNA is, in principle, a potent agonist of cytosolic RNA sensors such as RIG-I and MDA5, whose engagement normally triggers a signaling cascade through MAVS, TBK1/IKKε, and IRF3/IRF7 that culminates in type I and type III interferon production and the subsequent expression of hundreds of interferon-stimulated genes (ISGs). If the N protein interferes with this pathway, it could effectively delay or blunt the antiviral transcriptional program during the critical first hours of viral infection. This would allow incoming virions to establish replication complexes before the cell mounts a full defense, a strategy consistent with the observed early interferon suppression [68, 74]and delayed innate immune activation characteristic of SARS-CoV-2 infection [75].

An alternative, and not mutually exclusive, possibility is that N protein actively remodels the intracellular environment to generate a cellular state broadly permissive to viral replication. N protein is known to bind RNA promiscuously and to undergo liquid–liquid phase separation [33, 34]. Through such activities, N could alter stress granule dynamics, modulate protein translation, or reorganize membrane compartments that favor viral RNA replication. In this scenario, N protein functions less as a simple innate immune antagonist and more as a general reprogramming factor that shifts host cell physiology toward conditions optimized for coronavirus replication.

Equally compelling but less explored is the possibility that N protein neutralizes the antiviral activities of pre-existing or rapidly induced host restriction factors. Human cells constitutively express, or rapidly induce upon infection, a repertoire of restriction factors such as MX1, MX2, IFITM proteins, GADD34, APOBEC3 family members, TRIM5α, and BST2/tetherin that act at discrete steps of the viral life cycle[47, 76, 77]. It is reasonable to speculate that N protein evolved to counteract one or more of these restriction factors. Coronavirus RdRp complexes are essential for viral RNA replication and are thus vulnerable targets for host antiviral factors. Because N protein coats the viral genome and is present throughout the replication cycle, it is uniquely positioned to shield viral RNA and ribonucleoprotein complexes from exposure to these sensors and effectors.

In summary, the N protein of SARS-CoV-2 may promote infection through three overlapping mechanisms: suppressing RLR-dependent antiviral induction to benefit incoming virus, establishing an intracellular milieu intrinsically favorable to viral replication, and counteracting host restriction factors that would otherwise limit infection. Dissecting which of these mechanisms dominates, at which stages of the viral life cycle, and through which molecular interactions, would not only deepen our understanding of coronavirus pathogenesis but may also reveal new therapeutic targets.

## MATERIALS AND METHODS

### Plasmid construction and transfection

The pLVX-hACE2-hygro, pLVX-Flag-SARS-CoV-2 N, and SARS-CoV-2 GFP/ΔN (Wuhan-Hu-1, MN908947, Clade: 19A) [64] were kindly provided by Qiang Ding (Tsinghua University, Beijing). The cDNAs of SARS-CoV-2 N, SARS-CoV N, OC43-N, HKU1 N were synthesized with a C-terminal HA tag, cloned and inserted into the VR1012 vector by GentleGen (Suzhou, China). IRF3, ISRE, IFN-β, NF-κB luciferase reporter plasmids and Myc-cGAS, STING-Flag, MAVS-Flag, TRIF-Flag, MYD88-Flag, IKKβ-Flag, p65-Flag, TBK1-Flag, IRF3-Flag, MDA5 CARD-Myc, RIG-I CARD-Myc and HA-GADD34 expression plasmids were obtained as previously described [78-80].

The amino acid truncations SARS-CoV-2 N-HA 1–364,1-246, and 1-180 were synthesized with a C-terminal HA tag, cloned and inserted into the VR1012 vector by GentleGen (Suzhou, China). The plasmids were verified by sequencing before transfection using Hief Trans Liposomal Transfection Reagent (Yeasen Biotech, CN) following the manufacturer’s instructions.

### Antibodies and reagents

The antibodies and reagents used here are listed in table S1.

### Cell culture

HEK293T cells [CRL-3216, American Type Culture Collection (ATCC)] were cultured in Dulbecco’s modified Eagle’s medium supplemented with 10% fetal bovine serum (FBS) and 1% penicillin-streptomycin solution.

Caco-2 (HTB-37, ATCC) were cultured in RPMI 1640 medium supplemented with 10% FBS and 1% penicillin-streptomycin solution.

### Virus infection

For SeV, VSV and SARS-CoV-2 GFP/ΔN trVLPs, OC43 infection, HEK293T cells were transfected indicated plasmids for 18 hours and then infected with indicated virus. Caco-2 cells were infected with indicated virus. Eighteen hours later, the cells were collected, and viral infectivity was determined by fluorescence-activated cell sorting (FACS) or western blotting. GAPDH was used as a control.

### Stable expression cell line construction

Lentiviral plasmids were packaged into a lentivirus for infection of Caco-2 and HEK293T cells. Caco-2-N, HEK293T-ACE2 and HEK293T-ACE2-N stable expression cell lines were obtained under antibiotics selection.

### Immunoblotting and coimmunoprecipitation

Cell samples were harvested, washed twice with cold PBS, lysed in lysis buffer [50 mM tris (pH 7.5), with 150 mM NaCl, 0.5% NP-40, 0.5% deoxycholate sodium, 9 mM EDTA, and complete protease inhibitor cocktail tablets (Roche, USA)] at 4°C for 30 min, and then mixed into 1× loading buffer [0.08 M tris (pH 6.8), with 2.0% SDS, 10% glycerol, 0.1 M dithiothreitol, and 0.2% bromophenol blue] and boiled for 20 min. The cell lysates were separated on SDS-polyacrylamide gel electrophoresis (SDS-PAGE) gels and transferred to polyvinylidene fluoride membranes using a semidry apparatus (Bio-Rad, USA). Membranes were probed overnight with various primary antibodies against the proteins of interest. The secondary antibodies were horseradish peroxidase–conjugated anti-rabbit immunoglobulin G (IgG) (HuaBio, CN) and anti-mouse IgG (HuaBio, CN).

HEK293T cells were seeded into 6-cm dishes and transfected with a total of 3 μg of empty plasmid or various expression plasmids using Hief Trans Liposomal Transfection Reagent (Yeasen Biotech, CN). 24 h after transfection, the cells were lysed in lysis buffer [50 mM tris (pH 7.5), with 150 mM NaCl, 0.5% NP-40, and complete protease inhibitor cocktail tablets (Roche, USA)] at 4°C for 30 min. After sonication and centrifugation, the cell lysates were incubated with anti-Flag affinity matrix (Sigma-Aldrich, USA) or anti-HA affinity matrix (Roche, USA) at 4°C overnight on a rotator.

Staining was performed with enhanced chemiluminescence (ECL) substrate (Proteintech, USA) or Super ECL Detection Reagent chemiluminescent substrates (YeasenBiotech, CN) according to the manufacturer’s protocol. Membranes were imaged with a Series c300 (Azure Biosystems, USA).

### RNA extraction and qPCR

Total RNA from cells was isolated using RNA-Quick Purification Kit (RN001) and reverse-transcribed using Evo M-MLV RT Premix (AG11706) according to the manufacturer’s instructions. The complementary DNA was either undiluted or serially diluted in diethyl pyrocarbonate–treated water before real-time PCR to ensure that the amplification was within the linear range of detection. A SYBR Green Premix Kit (AG11701) was used for the RT-qPCR amplifications. The reactions were performed under the following conditions: 95°C for 30 s, followed by 40 cycles of 95°C for 5 s and 60°C for 34 s, and then a dissociation protocol. The presence of single peaks in the melting curve analysis indicated specific amplicons. The target primer sequences are provided in table S2.

### Dual luciferase assays

HEK293T cells were plated onto 24-well dishes and 100 ng of the firefly luciferase reporter plasmid as indicated, 5 ng of Renilla luciferase control plasmid (pRL-TK), and the indicated amounts of the expression plasmids were transfected per well. At 18 hours after transfection, the transactivation activity of the promoter was determined by measuring Firefly and Renilla luciferase activities using a Dual-Luciferase Reporter Assay Kit (catalog no. E2810, Promega, USA). Protein expression was confirmed by immunoblotting. Firefly luciferase activity was normalized to Renilla luciferase activity.

### Flow cytometry

After VSV and SARS-CoV-2 GFP/ΔN trVLPs infection, the Caco-2 cells were washed with cold PBS and analyzed on a DxFLEX flow cytometer (Beckman Coulter, USA). Data analysis was performed using FlowJo software.

### Phylogenetic Analysis

Nucleocapsid (N) protein amino acid sequences were retrieved from NCBI GenBank for 36 coronaviruses spanning four genera (α-CoV, β-CoV, γ-CoV, and δ-CoV). The dataset comprised 10 α-CoV, 3 β-CoV lineage A (Embecovirus), 12 β-CoV lineage B (Sarbecovirus), 3 β-CoV lineage C (Merbecovirus), 1 β-CoV lineage D (Nobecovirus), 2 γ-CoV, and 5 δ-CoV sequences. Accession numbers for all sequences are listed in Supplementary Table S3.

Amino acid sequences were aligned using MAFFT (L-INS-i strategy; -- localpair --maxiterate 1000) [81] as implemented in the NGPhylogeny.fr pipeline[82]. The resulting multiple sequence alignment was used for maximum-likelihood phylogenetic tree inference with IQ-TREE 2 [83]. The best-fit substitution model was automatically selected by ModelFinder [84] according to the Bayesian information criterion (BIC). Tree inference was performed under the selected model with gamma-distributed rate variation across sites. Branch support was assessed by 1,000 ultrafast bootstrap replicates [85]. The tree was rendered as an unrooted radial phylogeny.

### Prediction of N–MDA5 complexes

Complexes between SARS-CoV-2 N protein (full-length, residues 1–419, or the CTD alone, residues 247–365; GenBank YP_009724397.2, Wuhan-Hu-1) and the human MDA5 tandem CARDs (residues 1–200; UniProt Q9BYX4) were predicted using Boltz-2 with the managed MSA server for multiple-sequence alignment generation, using default inference parameters. For each prediction we recorded the aggregate confidence score, pLDDT, ipTM, pTM and predicted distance error metrics. Interface residues were defined as residue pairs with any heavy-atom distance below 5 Å between chains. Hydrogen bonds were assigned at donor–acceptor distances below 3.5 Å, salt bridges below 4.0 Å, and hydrophobic contacts below 5.0 Å. The predicted N-CTD binding surface was compared with the MDA5–MAVS CARD interface in the cryo-EM structure PDB 7DNI.

Chai-1 cross-validation. The same N-CTD–MDA5 CARDs complex was independently predicted with Chai-1[86] using the managed MSA server, with five diffusion samples. Interface residues and hydrogen bonds were defined identically as for Boltz-2. All five Chai-1 models were free of inter-chain clashes (ipTM 0.20–0.28). The MDA5-binding surface (CARD1 amino-terminal helix and CARD1–CARD2 linker) was reproduced in all five models, with MDA5 Tyr13 present at the interface in every model; the specific N-CTD contact patch differed between methods (Boltz-2, residues 279–281, 317–323 and 332–338; Chai-1, residues 338–364), and the Boltz-2-specific Thr332/Tyr333–Tyr13 hydrogen bonds were therefore not treated as independently confirmed.

### Statistical analysis

Data analyses were performed using GraphPad Prism 10.0 software. All data are shown as the means ± s.d.. The statistical significance analyses were performed using a two-sided unpaired t test. P values are indicated as follows: ns, not significant; *P < 0.05; **P < 0.01; ***P < 0.001.

## Acknowledgments

We thank Q. Dong for helping with the confocal microscopy analysis, M. Lou, M. Yu and W. Zheng for providing technical assistance, and S. Cen for providing OC43 viruses.

## Author contributions

J.S., Y.R., and Y.W. carried out most of the biochemical experiments with the help of S. C, Y. Cao, Y. Hu, S. Shen, and contributed to the supervision and data analysis. Q. Ding provided the key SARS-CoV-2 replicon viruses. X.-F.Y. directed the project, analyzed the data, and wrote the paper, with help from all of the authors.

## Data and materials availability

All data needed to evaluate the conclusions in the paper are present in the paper or the Supplementary Materials. All materials can be shared through a materials transfer agreement.

## References

1. Tanneti, N.S., H.A. Stillwell, and S.R. Weiss, Human coronaviruses: activation and antagonism of innate immune responses. Microbiol Mol Biol Rev, 2025. 89(1): p. e0001623.

2. Hoenigsperger, H., R. Sivarajan, and K.M. Sparrer, Differences and similarities between innate immune evasion strategies of human coronaviruses. Curr Opin Microbiol, 2024. 79: p. 102466.

3. Kasuga, Y., et al., Innate immune sensing of coronavirus and viral evasion strategies. Exp Mol Med, 2021. 53(5): p. 723–736.

4. Li, J.Y., et al., Innate Immunity Evasion Strategies of Highly Pathogenic Coronaviruses: SARS-CoV, MERS-CoV, and SARS-CoV-2. Front Microbiol, 2021. 12: p. 770656.

5. Otter, C.J., et al., Infection of primary nasal epithelial cells differentiates among lethal and seasonal human coronaviruses. Proc Natl Acad Sci U S A, 2023. 120(15): p. e2218083120.

6. Minkoff, J.M. and B. tenOever, Innate immune evasion strategies of SARS-CoV-2. Nat Rev Microbiol, 2023. 21(3): p. 178–194.

7. Nelemans, T. and M. Kikkert, Viral Innate Immune Evasion and the Pathogenesis of Emerging RNA Virus Infections. Viruses, 2019. 11(10).

8. Thorne, L.G., et al., Evolution of enhanced innate immune evasion by SARS-CoV-2. Nature, 2022. 602(7897): p. 487-495.

9. Rehwinkel, J. and M.U. Gack, RIG-I-like receptors: their regulation and roles in RNA sensing. Nat Rev Immunol, 2020. 20(9): p. 537–551.

10. Luan, X., et al., Innate immune responses to RNA: sensing and signaling. Front Immunol, 2024. 15: p. 1287940.

11. Thoresen, D., et al., The molecular mechanism of RIG-I activation and signaling. Immunol Rev, 2021. 304(1): p. 154–168.

12. Sharma, A., K. Kontodimas, and M. Bosmann, The MAVS Immune Recognition Pathway in Viral Infection and Sepsis. Antioxid Redox Signal, 2021. 35(16): p. 1376–1392.

13. Takaoka, A. and T. Yamada, Regulation of signaling mediated by nucleic acid sensors for innate interferon-mediated responses during viral infection. Int Immunol, 2019. 31(8): p. 477–488.

14. Yin, X., et al., MDA5 Governs the Innate Immune Response to SARS-CoV-2 in Lung Epithelial Cells. Cell Rep, 2021. 34(2): p. 108628.

15. Sampaio, N.G., et al., The RNA sensor MDA5 detects SARS-CoV-2 infection. Sci Rep, 2021. 11(1): p. 13638.

16. Thorne, L.G., et al., SARS-CoV-2 sensing by RIG-I and MDA5 links epithelial infection to macrophage inflammation. Embo j, 2021. 40(15): p. e107826.

17. Sampaio, N.G., et al., Loss of cellular RNA homeostasis contributes to MDA5 activation during virus infection. Nat Immunol, 2026. 27(9): p. 1815–1828.

18. Sánchez-Aparicio, M.T., et al., Paramyxovirus V Proteins Interact with the RIG-I/TRIM25 Regulatory Complex and Inhibit RIG-I Signaling. J Virol, 2018. 92(6).

19. Mandhana, R., L.K. Qian, and C.M. Horvath, Constitutively Active MDA5 Proteins Are Inhibited by Paramyxovirus V Proteins. J Interferon Cytokine Res, 2018. 38(8): p. 319–332.

20. Riedl, W., et al., Zika Virus NS3 Mimics a Cellular 14-3-3-Binding Motif to Antagonize RIG-I- and MDA5-Mediated Innate Immunity. Cell Host Microbe, 2019. 26(4): p. 493–503.e6.

21. Pulido, M.R., et al., MDA5 cleavage by the Leader protease of foot-and-mouth disease virus reveals its pleiotropic effect against the host antiviral response. Cell Death Dis, 2020. 11(8): p. 718.

22. Kim, H., et al., Foot-and-Mouth Disease Virus Evades Innate Immune Response by 3C-Targeting of MDA5. Cells, 2021. 10(2).

23. Liu, G., et al., ISG15-dependent activation of the sensor MDA5 is antagonized by the SARS-CoV-2 papain-like protease to evade host innate immunity. Nat Microbiol, 2021. 6(4): p. 467–478.

24. Sun, X., et al., Direct Interaction of Coronavirus Nonstructural Protein 3 with Melanoma Differentiation-Associated Gene 5 Modulates Type I Interferon Response during Coronavirus Infection. Int J Mol Sci, 2022. 23(19).

25. Zhang, X., et al., SARS-CoV-2 Nsp8 suppresses MDA5 antiviral immune responses by impairing TRIM4-mediated K63-linked polyubiquitination. PLoS Pathog, 2023. 19(11): p. e1011792.

26. Zheng, Y., et al., *Severe acute respiratory syndrome coronavirus 2 (SARS-CoV-2)* membrane *(M) protein inhibits type I and III interferon production by targeting RIG-I/MDA-5 signaling*. Signal Transduct Target Ther, 2020. 5(1): p. 299.

27. Han, L., et al., SARS-CoV-2 ORF9b antagonizes type I and III interferons by targeting multiple components of the RIG-I/MDA-5-MAVS, TLR3-TRIF, and cGAS-STING signaling pathways. J Med Virol, 2021. 93(9): p. 5376–5389.

28. Xia, H., et al., Evasion of Type I Interferon by SARS-CoV-2. Cell Rep, 2020. 33(1): p. 108234.

29. Hayn, M., et al., Systematic functional analysis of SARS-CoV-2 proteins uncovers viral innate immune antagonists and remaining vulnerabilities. Cell Rep, 2021. 35(7): p. 109126.

30. Cubuk, J., et al., The SARS-CoV-2 nucleocapsid protein is dynamic, disordered, and phase separates with RNA. Nat Commun, 2021. 12(1): p. 1936.

31. Dinesh, D.C., et al., Structural basis of RNA recognition by the SARS-CoV-2 nucleocapsid phosphoprotein. PLoS Pathog, 2020. 16(12): p. e1009100.

32. Carlson, C.R., et al., Phosphoregulation of Phase Separation by the SARS-CoV-2 N Protein Suggests a Biophysical Basis for its Dual Functions. Mol Cell, 2020. 80(6): p. 1092–1103.e4.

33. Perdikari, T.M., et al., SARS-CoV-2 nucleocapsid protein phase-separates with RNA and with human hnRNPs. Embo j, 2020. 39(24): p. e106478.

34. Savastano, A., et al., Nucleocapsid protein of SARS-CoV-2 phase separates into RNA-rich polymerase-containing condensates. Nat Commun, 2020. 11(1): p. 6041.

35. Oh, S.J. and O.S. Shin, SARS-CoV-2 Nucleocapsid Protein Targets RIG-I-Like Receptor Pathways to Inhibit the Induction of Interferon Response. Cells, 2021. 10(3).

36. Gori Savellini, G., et al., SARS-CoV-2 N Protein Targets TRIM25-Mediated RIG-I Activation to Suppress Innate Immunity. Viruses, 2021. 13(8).

37. Mu, J., et al., SARS-CoV-2 N protein antagonizes type I interferon signaling by suppressing phosphorylation and nuclear translocation of STAT1 and STAT2. Cell Discov, 2020. 6: p. 65.

38. Huang, C., et al., The Interaction between SARS-CoV-2 Nucleocapsid Protein and UBC9 Inhibits MAVS Ubiquitination by Enhancing Its SUMOylation. Viruses, 2023. 15(12).

39. Rui, Y., et al., Unique and complementary suppression of cGAS-STING and RNA sensing-triggered innate immune responses by SARS-CoV-2 proteins. Signal Transduct Target Ther, 2021. 6(1): p. 123.

40. Yoneyama, M., et al., The RNA helicase RIG-I has an essential function in double-stranded RNA-induced innate antiviral responses. Nat Immunol, 2004. 5(7): p. 730–7.

41. Thanos, D. and T. Maniatis, The high mobility group protein HMG I(Y) is required for NF-kappa B-dependent virus induction of the human IFN-beta gene. Cell, 1992. 71(5): p. 777–89.

42. Maniatis, T., et al., Structure and function of the interferon-beta enhanceosome. Cold Spring Harb Symp Quant Biol, 1998. 63: p. 609–20.

43. Wathelet, M.G., et al., Virus infection induces the assembly of coordinately activated transcription factors on the IFN-beta enhancer in vivo. Mol Cell, 1998. 1(4): p. 507–18.

44. Kawai, T. and S. Akira, Toll-like receptor and RIG-I-like receptor signaling. Ann N Y Acad Sci, 2008. 1143: p. 1–20.

45. Yoneyama, M., H. Kato, and T. Fujita, Physiological functions of RIG-I-like receptors. Immunity, 2024. 57(4): p. 731–751.

46. Wang, W., et al., Convergent Transcription of Interferon-stimulated Genes by TNF-α and IFN-α Augments Antiviral Activity against HCV and HEV. Sci Rep, 2016. 6: p. 25482.

47. Ye, R., et al., STING-NF-κB signaling builds an influenza spillover barrier. Science, 2026. 391(6788): p. eads4405.

48. Seth, R.B., et al., Identification and characterization of MAVS, a mitochondrial antiviral signaling protein that activates NF-kappaB and IRF 3. Cell, 2005. 122(5): p. 669–82.

49. Zandi, E., et al., The IkappaB kinase complex (IKK) contains two kinase subunits, IKKalpha and IKKbeta, necessary for IkappaB phosphorylation and NF-kappaB activation. Cell, 1997. 91(2): p. 243–52.

50. Schmitz, M.L. and P.A. Baeuerle, The p65 subunit is responsible for the strong transcription activating potential of NF-kappa B. Embo j, 1991. 10(12): p. 3805–17.

51. Fitzgerald, K.A., et al., IKKepsilon and TBK1 are essential components of the IRF3 signaling pathway. Nat Immunol, 2003. 4(5): p. 491–6.

52. Yoneyama, M., et al., Shared and unique functions of the DExD/H-box helicases RIG-I, MDA5, and LGP2 in antiviral innate immunity. J Immunol, 2005. 175(5): p. 2851–8.

53. Liu, S., et al., Phosphorylation of innate immune adaptor proteins MAVS, STING, and TRIF induces IRF3 activation. Science, 2015. 347(6227): p. aaa2630.

54. Song, B., et al., Ordered assembly of the cytosolic RNA-sensing MDA5-MAVS signaling complex via binding to unanchored K63-linked poly-ubiquitin chains. Immunity, 2021. 54(10): p. 2218–2230.e5.

55. Cai, X., et al., Prion-like polymerization underlies signal transduction in antiviral immune defense and inflammasome activation. Cell, 2014. 156(6): p. 1207–1222.

56. Wu, B., et al., Molecular imprinting as a signal-activation mechanism of the viral RNA sensor RIG-I. Mol Cell, 2014. 55(4): p. 511–23.

57. Timani, K.A., et al., Nuclear/nucleolar localization properties of C-terminal nucleocapsid protein of SARS coronavirus. Virus Res, 2005. 114(1-2): p. 23–34.

58. Wang, M., M. Valadez-Ingersoll, and T.D. Gilmore, Control of nuclear localization of the nucleocapsid protein of SARS-CoV-2. Virology, 2024. 600: p. 110232.

59. Passaro, S., et al., Boltz-2: Towards Accurate and Efficient Binding Affinity Prediction. 2025: p. 2025.06.14.659707.

60. Evans, R., et al., Protein complex prediction with AlphaFold-Multimer. 2022: p. 2021.10.04.463034.

61. Motz, C., et al., Paramyxovirus V proteins disrupt the fold of the RNA sensor MDA5 to inhibit antiviral signaling. Science, 2013. 339(6120): p. 690–3.

62. McBride, R., M. van Zyl, and B.C. Fielding, The coronavirus nucleocapsid is a multifunctional protein. Viruses, 2014. 6(8): p. 2991–3018.

63. Cong, Y., et al., Nucleocapsid Protein Recruitment to Replication-Transcription Complexes Plays a Crucial Role in Coronaviral Life Cycle. J Virol, 2020. 94(4).

64. Ju, X., et al., A novel cell culture system modeling the SARS-CoV-2 life cycle. PLoS Pathog, 2021. 17(3): p. e1009439.

65. Otter, C.J., et al., Interferon signaling in the nasal epithelium distinguishes among lethal and common cold coronaviruses and mediates viral clearance. Proc Natl Acad Sci U S A, 2024. 121(21): p. e2402540121.

66. Hadjadj, J., et al., Impaired type I interferon activity and inflammatory responses in severe COVID-19 patients. Science, 2020. 369(6504): p. 718–724.

67. Lucas, C., et al., Longitudinal analyses reveal immunological misfiring in severe COVID-19. Nature, 2020. 584(7821): p. 463–469.

68. Blanco-Melo, D., et al., Imbalanced Host Response to SARS-CoV-2 Drives Development of COVID-19. Cell, 2020. 181(5): p. 1036–1045.e9.

69. Pichlmair, A., et al., RIG-I-mediated antiviral responses to single-stranded RNA bearing 5’-phosphates. Science, 2006. 314(5801): p. 997–1001.

70. Kato, H., et al., Differential roles of MDA5 and RIG-I helicases in the recognition of RNA viruses. Nature, 2006. 441(7089): p. 101–5.

71. Knoops, K., et al., SARS-coronavirus replication is supported by a reticulovesicular network of modified endoplasmic reticulum. PLoS Biol, 2008. 6(9): p. e226.

72. Li, Y., et al., SARS-CoV-2 induces double-stranded RNA-mediated innate immune responses in respiratory epithelial-derived cells and cardiomyocytes. Proc Natl Acad Sci U S A, 2021. 118(16).

73. Yao, H., et al., Molecular Architecture of the SARS-CoV-2 Virus. Cell, 2020. 183(3): p. 730–738.e13.

74. V’Kovski, P., et al., Coronavirus biology and replication: implications for SARS-CoV-2. Nat Rev Microbiol, 2021. 19(3): p. 155–170.

75. Perlman, S. and J. Netland, Coronaviruses post-SARS: update on replication and pathogenesis. Nat Rev Microbiol, 2009. 7(6): p. 439–50.

76. Kluge, S.F., D. Sauter, and F. Kirchhoff, SnapShot: antiviral restriction factors. Cell, 2015. 163(3): p. 774–774.e1.

77. Ishaq, M., H. Marshall, and V. Natarajan, GADD34 attenuates HIV-1 replication by viral 5’-UTR TAR RNA-mediated translational inhibition. Virology, 2020. 540: p. 119–131.

78. Su, J., et al., HIV-2/SIV Vpx targets a novel functional domain of STING to selectively inhibit cGAS-STING-mediated NF-κB signalling. Nat Microbiol, 2019. 4(12): p. 2552–2564.

79. Rui, Y., et al., Disruption of MDA5-Mediated Innate Immune Responses by the 3C Proteins of Coxsackievirus A16, Coxsackievirus A6, and Enterovirus D68. J Virol, 2017. 91(13).

80. Rui, Y., et al., HIV-1 Vpu and SARS-CoV-2 ORF3a proteins disrupt STING-mediated activation of antiviral NF-κB signaling. Sci Signal, 2025. 18(870): p. eadd6593.

81. Katoh, K. and D.M. Standley, MAFFT multiple sequence alignment software version 7: improvements in performance and usability. Mol Biol Evol, 2013. 30(4): p. 772–80.

82. Lemoine, F., et al., NGPhylogeny.fr: new generation phylogenetic services for non-specialists. Nucleic Acids Res, 2019. 47(W1): p. W260–w265.

83. Minh, B.Q., et al., IQ-TREE 2: New Models and Efficient Methods for Phylogenetic Inference in the Genomic Era. Mol Biol Evol, 2020. 37(5): p. 1530–1534.

84. Kalyaanamoorthy, S., et al., ModelFinder: fast model selection for accurate phylogenetic estimates. Nat Methods, 2017. 14(6): p. 587–589.

85. Hoang, D.T., et al., UFBoot2: Improving the Ultrafast Bootstrap Approximation. Mol Biol Evol, 2018. 35(2): p. 518–522.

86. Discovery, C., et al., Chai-1: Decoding the molecular interactions of life. 2024: p. 2024.10.10.615955.

