## Supplementary material for "Coronavirus nucleocapsid proteins differentially antagonize RIG-I-like receptor signaling to shape pathogenicity": sup figures

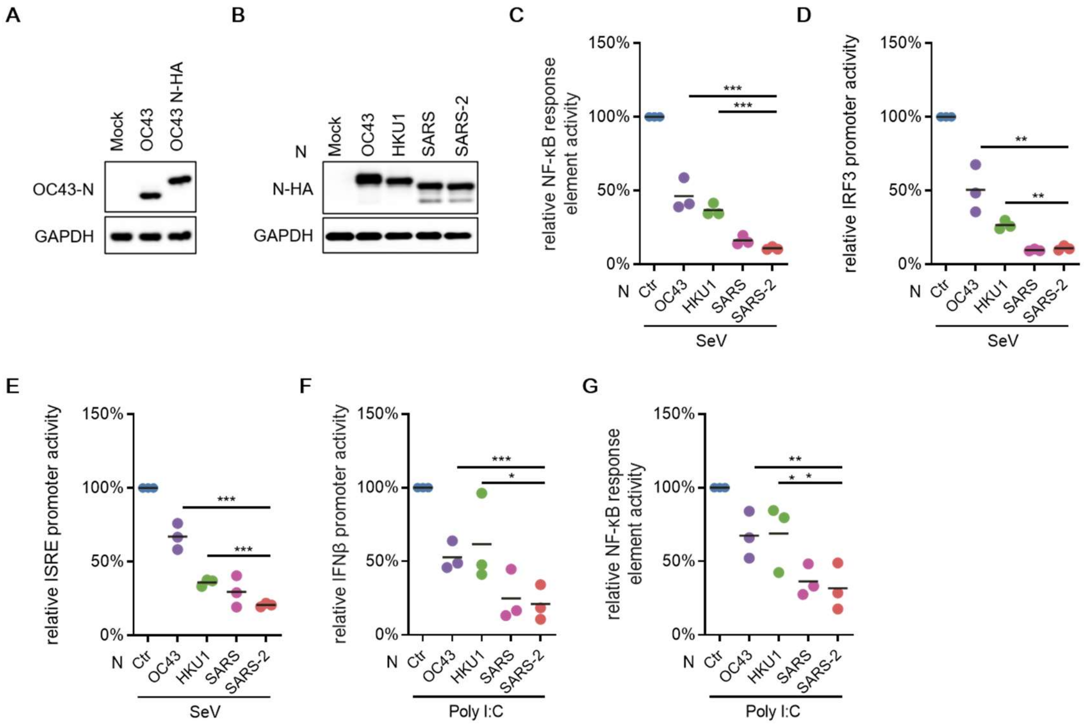


**Fig. S1 | N proteins from highly pathogenic human coronaviruses suppress SeV- and poly (I:C)-induced innate immune activation.**

**A,** HEK293T cells were infected with HCoV-OC43 at an MOI of 0.5, or mock-infected, together with transfection of OC43 N-HA (200 ng); 18 h later, cell lysates were immunoblotted for viral N. **B,** HEK293T cells were transfected HA-tagged N from the indicated viruses, 18 h later, cell lysates were immunoblotted for HA antibody. **C,** HEK293T cells were co-transfected with an NF-κB response element reporter, a Renilla internal control and HA-tagged N from the indicated viruses; 18 h after transfection, cells were stimulated with SeV and harvested 18 h later. **D,E,** As in **C**, using an IRF3 promoter reporter (**D**) or ISRE reporter (**E**). **F,G,** HEK293T cells were co-transfected with an IFNβ promoter (**F**) or NF-κB response element (**G**) reporter, a Renilla internal control, poly (I:C) and HA-tagged N from the indicated viruses, and harvested 36 h after transfection. Reporter activity in stimulated cells lacking N was set to 100%. Data are mean ± s.d. of three independent biological replicates; P values were calculated by two-sided unpaired t-tests comparing the indicated N protein with SARS-CoV-2 N (***$P < 0.001$, **$P < 0.01$, *$P < 0.05$).

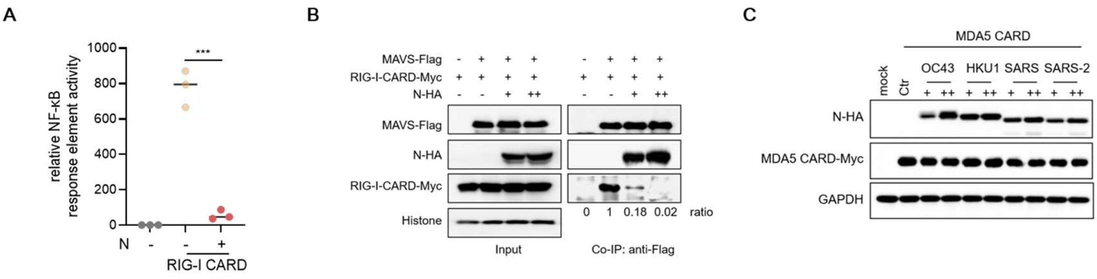


**Fig. S3 | The SARS-CoV-2 N protein interacts with the CARD domains of RIG-I.**

**A,** HEK293T cells were co-transfected with an NF-κB response element reporter, a Renilla internal control, RIG-I CARD and increasing doses of SARS-CoV-2-N-HA as indicated, and harvested 36 h after transfection. **B,** HEK293T cells were co-transfected with MAVS-Flag, RIG-I CARD-Myc and increasing doses of SARS-CoV-2-N-HA as indicated; 36 h later, cell lysates were subjected to anti-Flag IP followed by immunoblot analysis of co-precipitated RIG-I CARD-Myc, N-HA. **C,** immunoblot analysis corresponding to Fig. 3d. Data are mean ± s.d. of three independent biological replicates; P values were calculated by two-sided unpaired t-tests comparing each condition with RIG-I CARD alone (***P < 0.001).

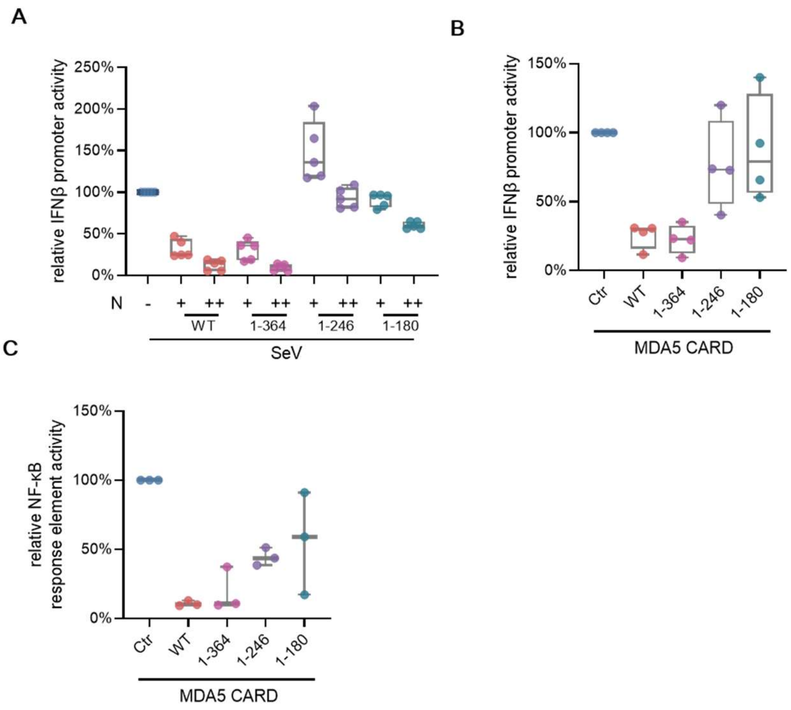


**Fig. S4 | The CTD mediates N suppression of IFNβ and NF-κB signalling.**

**A,** HEK293T cells were co-transfected with an IFNβ promoter reporter, a Renilla internal control and the indicated HA-tagged N truncation mutants; 18 h after transfection, cells were stimulated with SeV and harvested 18 h later. Reporter activity in SeV-stimulated cells was set to 100%. **B,C,** HEK293T cells were co-transfected with an IFNβ promoter (**B**) or NF-κB response element (**C**) reporter, a Renilla internal control, MDA5 CARD and the indicated N truncation mutants, and harvested 36 h after transfection; reporter activity in cells expressing MDA5 CARD alone was set to 100%. Data are mean ± s.d. of three independent biological replicates.

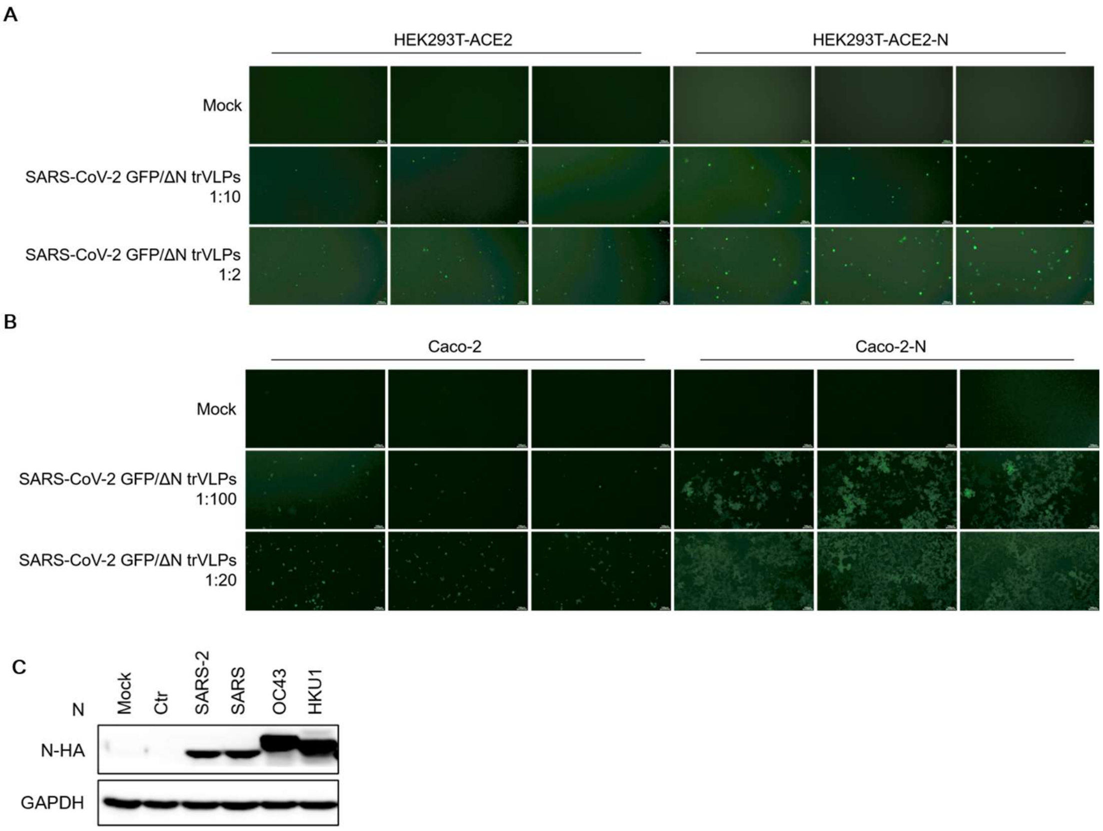


Fig. S5 | GFP imaging of SARS-CoV-2 replicon-infected cells corresponding to Fig. 5b,c.

A-B, Representative GFP fluorescence images and WB of the samples analysed in Fig. 5B,C; C, WB of the samples in Fig.5D

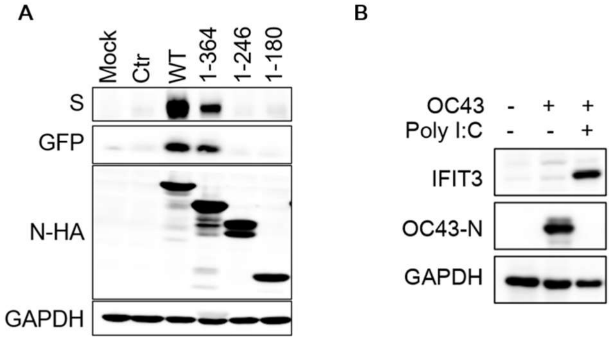


Fig. S6 RLR activation suppresses OC43 replication

A, Representative WB of the samples analysed in Fig. 6A; B, HEK239T cells were transfected with Poly (I:C) for 18 h, then infected with OC43 viruses; 18 h later, cell lysates were immunoblotted for viral N protein andIFIT3, with GAPDH as loading control.
